# Common and distinct contributions of the dorsomedial prefrontal and posterior parietal cortices to social episodic memory

**DOI:** 10.64898/2026.01.16.699750

**Authors:** C. M. Bates, A. K. Martin

## Abstract

The self-reference effect (SRE) refers to the robust mnemonic advantage for self-relevant information and is characterised by a graded pattern spanning the self, close others, and distant others. Despite extensive behavioural evidence, causal evidence for the neural mechanisms supporting the SRE remains limited, particularly during incidental encoding and across source memory, metacognition, and emotional valence. We investigated the causal contributions of the dorsomedial prefrontal cortex (dmPFC) and posterior parietal cortex (PPC) to self–other episodic memory and metacognition using focal transcranial direct current stimulation (tDCS). One hundred participants completed an incidental encoding task in which trait adjectives were rated with reference to the self, a close friend, or a celebrity, under anodal or sham stimulation to either region. Recognition tests assessed item and source memory sensitivity (d′) and metacognitive efficiency (meta-d′/d′) as a function of agent and emotional valence. Across stimulation conditions, a graded advantage for self-related information was observed in both item and source memory, extending to metacognitive efficiency. Anodal stimulation to either region attenuated the self-reference effect. PPC stimulation additionally biased source memory away from positive and towards negative information. These findings demonstrate shared and dissociable causal roles of dmPFC and PPC in self–other episodic memory.

---

Humans exhibit systematic self-biases in cognitive processing, one prominent example being the self-reference effect (SRE) in episodic memory. The SRE refers to the reliable enhancement of memory for self-relevant information relative to information encoded in other, non-self-referential contexts (Symons & Johnson, 1997). This advantage is commonly attributed to deeper and more elaborative encoding of self-relevant material, which benefits from integration with rich, well-established self-related knowledge structures (Conway, 2005). Importantly, the magnitude of the SRE is not categorical but graded as a function of social closeness. Memory performance is typically strongest for self-referenced information, intermediate for information related to close others, and weakest for information associated with socially distant individuals such as celebrities (Kokici et al., 2023). Despite substantial behavioural evidence for the self-reference effect, causal evidence for its neural substrates remains limited. Here, we used focal transcranial direct current stimulation targeting the dorsomedial prefrontal cortex and posterior parietal cortex to test their roles in graded self–other episodic memory.

The medial prefrontal cortex (mPFC) is a core region of the social brain implicated in the processing of both self- and other-relevant information (Courtney & Meyer, 2020; Denny et al., 2012), as well as in episodic memory encoding and retrieval (Euston et al., 2012). Neuroimaging studies suggest a ventral-to-dorsal functional gradient within the mPFC, whereby ventral regions are preferentially engaged during self-referential processing, while more dorsal regions are recruited during other-referential processing (Denny et al., 2012). Consistent with this account, causal evidence from non-invasive brain stimulation indicates that focal transcranial direct current stimulation to the dorsomedial prefrontal cortex (dmPFC) selectively enhances recognition for other-relevant, but not self-relevant, information (Martin, Huang, et al., 2019; Martin, Su, et al., 2019). However, these studies employed intentional encoding with an expected memory test, raising the possibility that executive control processes were disproportionately engaged, potentially inflating prefrontal contributions to memory performance (Kensinger & Ford, 2021). Moreover, prior stimulation work has focused primarily on the self and socially distant others, despite evidence that close others occupy a distinct and potentially intermediate representational space in the mPFC (Courtney & Meyer, 2020).

The posterior parietal cortex (PPC) is often associated with attentional orienting, perceptual processing, and sensorimotor integration (Corbetta & Shulman, 2011), but it is also increasingly recognised as a key contributor to episodic memory encoding, consolidation, and retrieval (Berryhill, 2012; Sestieri et al., 2017). Brain stimulation studies targeting the PPC have yielded mixed effects on memory performance, with some reports demonstrating modulation of associative memory (Meng et al., 2021; Vulić et al., 2021) and others observing null effects (Dubravac et al., 2021; Rossi et al., 2006). Notably, the medial parietal cortex, a subregion within the broader PPC, has been implicated in self- and other-referential memory retrieval, showing a graded pattern of activation that is strongest for the self, intermediate for close others, and weakest for public figures (Lou et al., 2004). Moreover, disruption of medial parietal activity using transcranial magnetic stimulation selectively impairs self-referential memory retrieval (Lou et al., 2004). Beyond episodic memory, converging evidence suggests that the PPC plays a broader role in representing social space, mapping relationships between the self and others in a manner analogous to its role in representing physical space (Yamakawa et al., 2009). Together, these findings suggest that the PPC contributes not only to memory processes but also to the self–other distinctions that underpin social cognition.

An important component that has received relatively little attention in research on the self-reference effect is metamemory, or metacognition about memory, defined as the ability to monitor and evaluate one’s own memory performance (Dunlosky & Thiede, 2013). Although self-biases are well established across core cognitive domains including memory, attention, and perception (Amodeo et al., 2021; Nijhof et al., 2020), it remains unclear whether these biases extend to higher-order processes such as metacognitive awareness of memory. Neurocognitive models of metacognition implicate distributed frontoparietal systems, including medial prefrontal and posterior parietal regions that overlap with networks supporting self-referential processing and episodic memory (Fleming & Dolan, 2012). Kokici and colleagues (2023) addressed this question by applying signal detection theoretic measures to examine whether the graded self–other pattern observed in memory performance is also evident in metamemory. Specifically, they quantified metacognitive sensitivity (meta-d′), which reflects how effectively confidence judgements distinguish between correct and incorrect memory decisions. Because meta-d′ is constrained by first-order memory sensitivity (d′), metacognitive efficiency, defined as the ratio of meta-d′ to d′, was also calculated to isolate metacognitive performance independently of memory accuracy (Fleming & Lau, 2014). Kokici and colleagues (2023) found that the characteristic graded self–other pattern was present in metacognitive sensitivity but absent in metacognitive efficiency. This dissociation suggests that self-biases in episodic memory may be supported by neural mechanisms that influence memory encoding or retrieval, rather than by enhanced efficiency of metacognitive monitoring.

A critical component of self- and other-referential episodic memory is the binding of contextual details, including information about the agent to whom a memory is attributed, a process commonly referred to as source monitoring. Source memory is often considered a more sensitive index of episodic memory quality than item memory alone, as it requires the integration of content with contextual features present at encoding (Leshikar & Duarte, 2012). Neurocognitive accounts of source monitoring implicate interactions between medial prefrontal regions involved in self- and other-referential processing and posterior parietal regions that support attentional orienting, contextual binding, and retrieval monitoring (Dobbins et al., 2002; Sestieri et al., 2017). Consistent with this framework, empirical evidence suggests that the self-reference effect extends beyond item recognition to influence source memory, indicating a qualitative enhancement of episodic representations. For example, self-referential encoding has been shown to improve both item and source recognition, consistent with more elaborative or distinctive memory traces (Durbin et al., 2017; Leshikar & Duarte, 2012; Serbun et al., 2011). However, there remains ongoing debate as to whether source memory reflects a distinct cognitive process or an extension of item memory supported by shared mechanisms (Chirtop et al., 2025; Davachi et al., 2003; Dent & Martin, 2023; Glisky et al., 1995; Guo et al., 2021; Mather, 2007; Slotnick et al., 2003). In the context of self-referential processing, resolving this issue is particularly important, as source memory may differentially depend on medial prefrontal and parietal contributions to self–other representation and episodic retrieval.

Finally, it is important to consider the role of emotional valence in the self-reference effect. Prior research has demonstrated a positivity bias in self-referential memory, whereby individuals preferentially remember positive information about themselves, but not necessarily about socially distant others (D’Argembeau et al., 2005). Neurocognitive accounts suggest that such valence-dependent self-biases may arise from interactions between medial prefrontal systems involved in self-evaluation and posterior parietal regions supporting attentional allocation and memory monitoring (Cabeza et al., 2008; Moran et al., 2006). More recently, Kokici and colleagues (2023) applied first- and second-order signal detection analyses to examine both memory sensitivity and metacognitive monitoring across self-, close-other, and public-figure encoding conditions. They observed a general positivity bias in recognition memory across all agents, with the magnitude of this bias graded as strongest for the self, intermediate for a close friend, and weakest for a celebrity. Importantly, although positive items were recognised with higher accuracy, they were associated with reduced metacognitive sensitivity across agents, suggesting a dissociation between memory strength and confidence calibration. In source memory, self-referential advantages were observed primarily for positive information and were driven by reduced accuracy in attributing positive traits to socially distant others. Together, these findings indicate that the influence of valence on memory and memory monitoring reflects the joint contribution of self-referential and attentional monitoring systems, and varies systematically with social closeness.

To our knowledge, no previous studies have used non-invasive brain stimulation to examine the self-reference effect under conditions of incidental encoding while simultaneously assessing source memory, metacognitive efficiency, and emotional valence. Although prior stimulation work has focused primarily on medial prefrontal contributions to intentional self-referential memory, far less is known about the causal role of posterior parietal regions, which are implicated in attentional control, source monitoring, and memory evaluation. Moreover, while emotional valence is known to modulate self-related memory biases, its interaction with self–other encoding and metamemory at the neural level remains largely unexplored.

Accordingly, the present study applied focal transcranial direct current stimulation to either the dorsomedial prefrontal cortex or the posterior parietal cortex during an incidental self-and other-referential encoding task involving trait judgements for the self, a close friend, and a celebrity. Subsequent recognition tests assessed both item and source memory, alongside metacognitive efficiency derived from signal detection theoretic models. By orthogonally manipulating social agent and emotional valence, this design allowed us to test the independent and shared causal contributions of medial prefrontal and parietal systems to graded self–other memory effects, memory monitoring, and valence-dependent biases.

## Methods

### Participants

One hundred participants were recruited via the University of Kent Research Participation Scheme and local advertisements, and received either course credit or monetary compensation for their time. Participants were randomly assigned to one of four between-subjects conditions (N = 25 per condition). All participants had normal or corrected-to-normal vision, reported no history of neurological or psychiatric conditions, and had no non-removable electrical medical devices. Ethical approval was obtained from the Psychology Ethics Committee at the University of Kent.

### Design

The study employed a 2 × 2 × 3 × 2 mixed-design analysis of variance, with two between-subjects factors (stimulation site and stimulation type) and two within-subjects factors (agent and emotional valence). An a priori power analysis conducted using G*Power (Faul et al., 2007) indicated that a sample size of 100 provided 80 percent power to detect medium-sized between-subjects effects (Cohen’s f approximately 0.25 to 0.29) and small-to-medium within-subjects effects and interactions (Cohen’s f approximately 0.13 to 0.14), assuming sphericity and an alpha level of .05.

### Materials

#### Hardware and Software

The experimental task was programmed and presented using PsychoPy3 (Peirce et al., 2019) and administered on a standard desktop computer with a monitor, mouse, and keyboard. Questionnaires and demographic information were completed using paper-based forms.

#### Measures

All participants completed a tDCS safety screening questionnaire to confirm eligibility for stimulation. Mood was assessed using the Visual Analogue Mood Scale (VAMS; Folstein & Luria, 1973) to evaluate changes in positive and negative affect following stimulation in both anodal and sham conditions. Adverse effects associated with stimulation were assessed using an adapted version of the questionnaire developed by Brunoni et al. (2011), administered to participants in all stimulation groups.

##### Self-Other Referential Memory Task

Episodic memory was assessed using a self–other referential paradigm in which trait adjectives were encoded with reference to the self, a close friend, or a celebrity, specifically Boris Johnson, who was the UK Prime Minister at the time of testing. During the encoding phase, participants rated each target on 20 trait adjectives using a nine-point scale ranging from very inaccurate to very accurate. Each target was paired with a unique set of adjectives selected from the normative database reported by Warriner et al. (2013) and matched across conditions for emotional valence and arousal. Stimuli were presented in a pseudorandom order, with no more than two consecutive trials from the same agent condition. Following encoding, participants completed an unrelated distractor task lasting approximately five minutes.

In the retrieval phase, participants completed a surprise recognition test in which the 60 previously encoded adjectives were intermixed with 60 valence- and arousal-matched distractor adjectives. For each word, participants made an old–new recognition judgement. For items endorsed as previously seen, participants then indicated the agent with which the adjective had been associated during encoding. Confidence ratings were collected for both item recognition and source memory judgements, allowing computation of memory sensitivity and metacognitive efficiency.

##### Focal tDCS

The focal tDCS setup included a single-channel direct current stimulator (DC-Stimulator Plus, NeuroConn), with a central electrode (2.5cm diameter) and return electrode (inner/outer diameter of 7.5cm and 9.8cm, respectively) attached using adhesive electroconductive gel and an EEG cap. The central electrode determines stimulation polarity, in this case anodal, and the return electrode constrains electrical current to the target region, preserving spatial focality (Gbadeyan et al., 2016; Villamar et al., 2013). During anodal sessions, transcranial stimulation was delivered at 1 mA for 20 minutes, with a 5-second ramp-up and ramp-down at the beginning and end of stimulation. In the sham condition, stimulation was applied for 40 seconds, including the same ramp-up and ramp-down procedure, to mimic the initial sensation of stimulation while avoiding any lasting neurophysiological effects. The dmPFC was located by finding 15% of the distance from the Fz towards the Fpz, whilst the PPC was located by finding Pz. Current modelling was completed using SimNIBS version 4.1 (Thielscher et al., 2015). Parameters were selected to estimate current delivery according to EEG 10-20 guided electrode positions as stated above, with the return electrode placed equidistantly around the center electrode. Electrode and gel thickness were modelled as 2 and 1mm, respectively. Standard conductivity values as provided by SimNIBS were used. We present the theoretical cortical electrical field (magnitude E) for anodal focal-tDCS to both the dmPFC and the PPC (see Figure 1).

**Figure 1.**
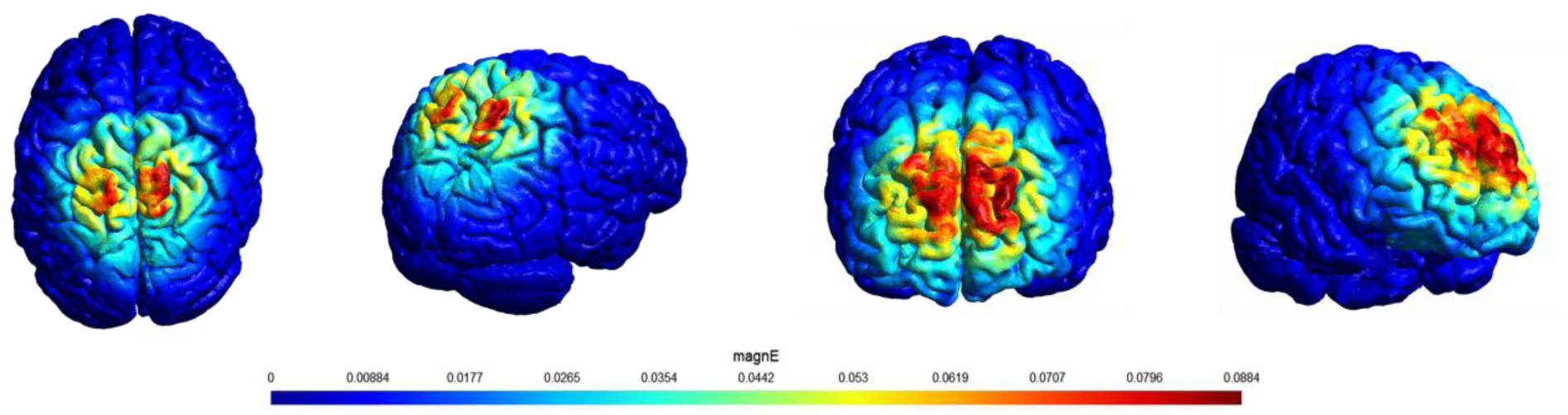
Focal-tDCS Current Modelling Across PPC and dmPFC Regions. *Note.* Figures depict the flow and spread of electrical current when delivered at 1mA to the PPC and the dmPFC, respectively.

### Procedure

Participants were first presented with information about the study and provided informed written consent for their participation. Participants completed a safety screening form, demographic questionnaire, and the VAMS. Stimulation was administered during the encoding phase of the memory task. Blinding was achieved by using the “study-mode” and a set of codes administered by a researcher not actively participating in data collection. Upon completion of the task, participants completed the VAMS and adverse effects checklist. To assess blinding, participants were asked to guess whether they received sham or anodal stimulation. Finally, participants were debriefed and compensated for their time.

### Statistical Analyses

Item memory performance was analysed using a 2×2×3×2 mixed-design analysis of variance (ANOVA). The between-subjects factors were Region (dorsomedial prefrontal cortex, posterior parietal cortex) and Stimulation Type (anodal, sham), and the within-subjects factors were Agent (self, close other, celebrity) and Valence (positive, negative). The dependent variable was discrimination sensitivity (d′), calculated according to signal detection theory as the difference between the z-transformed hit rate and false alarm rate.

Metacognitive performance was quantified following the signal detection theoretic framework described by Fleming (2017). For each participant, metacognitive sensitivity (meta-d′) and metacognitive efficiency (meta-d′/d′) were computed based on confidence ratings. Metacognitive efficiency was the primary index of interest and was entered as the dependent variable in separate 2×2×3×2 mixed-design ANOVAs with the same factors as above.

Source memory was analysed using an analogous approach. Source discrimination sensitivity (source d′) was entered as the dependent variable in a further set of 2×2×3×2 mixed-design ANOVAs. Source-related metacognitive efficiency (meta-d′/d′) was calculated from confidence ratings associated with source judgements and analysed using the same factorial structure.

To assess potential effects of stimulation on mood, composite scores for negative and positive affect were calculated at pre- and post-stimulation. Negative affect comprised ratings of afraid, confused, sad, angry, tired, and tense, whereas positive affect comprised ratings of energetic and happy. Change scores were computed by subtracting pre-stimulation from post-stimulation values and analysed using ANOVAs with Region and Stimulation Type as between-subjects factors. Adverse effects were quantified by summing responses across all items of the adverse effects questionnaire, yielding a total score for each participant, which was analysed using the same between-subjects ANOVA structure.

Blinding accuracy (correct vs. incorrect guesses) was analysed using contingency and binomial approaches. Differences in accuracy between regions were assessed using a chi-square test of independence. To characterise performance relative to chance (0.5) within each region, exact binomial tests were conducted separately for each region.

## Results

A 2 (Stimulation condition) × 2 (Target region) between-subjects ANOVA confirmed that groups were age-matched. There were no main effects of stimulation, *F*(1, 96) = 2.10, *p* = .15, η²_p_ = 0.02, or region, *F*(1, 96) = 0.65, *p* = .42, η²_p_ = 0.01, and no stimulation × region interaction, *F*(1, 96) = 0.03, *p* = .87, η²_p_ < 0.001. Descriptively, mean age ranged from 19.36 to 20.48 years across cells.

Sex distribution did not differ across experimental groups. Within each region, the distribution of sex across stimulation conditions was comparable (Region 1: χ²(1, N = 50) = 0.94, p = .33; Region 2: χ²(1, N = 50) = 0.10, p = .75), and collapsing across regions there was no overall association between sex and stimulation condition (χ²(1, N = 100) = 0.21, p = .65).

Only hypotheses relevant findings will be reported in-text. For all analysis outputs please refer to appendix 1.

### Item Memory

Means and standard deviations across all conditions are presented in Table 1.

**Table 1.** Means and standard deviations across all tasks as per stimulation condition and region of stimulation for item memory.

| Outcome | Valence | Agent | dmPFC – Sham | dmPFC – Anodal | PPC – Sham | PPC – Anodal |
| --- | --- | --- | --- | --- | --- | --- |
| Sensitivity ($d'$) | Positive | Self | 2.08 (0.60) | 1.87 (0.59) | 2.24 (0.82) | 2.06 (0.83) |
|  |  | Friend | 1.75 (0.46) | 1.71 (0.49) | 1.89 (0.64) | 1.78 (0.76) |
|  |  | Celeb | 1.14 (0.64) | 1.29 (0.47) | 1.24 (0.61) | 1.31 (0.67) |
|  | Negative | Self | 2.22 (0.38) | 2.03 (0.63) | 2.18 (0.67) | 2.01 (0.59) |
|  |  | Friend | 1.88 (0.60) | 1.73 (0.67) | 1.94 (0.69) | 1.83 (0.69) |
|  |  | Celeb | 1.36 (0.62) | 1.60 (0.68) | 1.25 (0.64) | 1.27 (0.53) |
| Metacognitive efficiency<br>(meta-$d'/d'$) | Positive | Self | 0.77 (0.34) | 0.77 (0.23) | 0.97 (0.46) | 0.83 (0.37) |
|  |  | Friend | 0.71 (0.24) | 0.75 (0.25) | 0.80 (0.42) | 0.63 (0.25) |
|  |  | Celeb | 0.43 (0.08) | 0.45 (0.07) | 0.55 (0.28) | 0.45 (0.23) |
|  | Negative | Self | 0.75 (0.21) | 0.76 (0.21) | 0.79 (0.41) | 0.61 (0.29) |
|  |  | Friend | 0.81 (0.41) | 0.79 (0.30) | 0.75 (0.29) | 0.61 (0.22) |
|  |  | Celeb | 0.69 (0.23) | 0.71 (0.21) | 0.55 (0.20) | 0.47 (0.10) |

#### Memory Sensitivity (d’)

A significant main effect of Agent was observed, *F*(2, 192) = 167.62, *p* < .001, η²G = 0.21. Memory sensitivity was highest for self-referenced items, intermediate for items encoded in relation to a close other, and lowest for items encoded in relation to a distant other. Post hoc comparisons confirmed that memory sensitivity differed significantly between all three agent conditions. Specifically, self-referenced items were remembered more accurately than items encoded in relation to a close other, *t*(99) = 3.59, *p* < .001, *d* = 0.43, and more accurately than items encoded in relation to a distant other, *t*(99) = 10.70, *p* < .001, *d* = 1.23. In addition, items encoded in relation to a close other were remembered more accurately than those encoded in relation to a distant other, *t*(99) = 6.92, *p* < .001, *d* = 0.80.

There was also a significant interaction between Stimulation Type and Agent, *F*(2, 192) = 6.82, *p* = .001, η²G = .01. Follow-up analyses examining the effect of stimulation separately within each agent condition revealed no statistically significant differences between anodal and sham stimulation for self-referenced items, *t*(98) = 1.81, *p* = .07, *d* = 0.36, close-other-referenced items, *t*(98) = 0.92, *p* = .36, *d* = 0.18, or distant-other-referenced items, *t*(98) = −1.20, *p* = .23, *d* = −0.24.

When the data were examined separately by stimulation condition, the effect of Agent remained significant under both sham and anodal stimulation. Under sham stimulation, a robust agent effect was observed, *F*(2, 98) = 99.26, *p* < .001, η²G = 0.38. Under anodal stimulation, the agent effect was still significant but reduced in magnitude, *F*(2, 98) = 66.89, *p* < .001, η²G = 0.14. This pattern indicates that stimulation to either the dorsomedial prefrontal cortex or the posterior parietal cortex attenuated the self-reference effect in item memory. Inspection of the means suggests that this reduction reflected a combination of decreased memory sensitivity for self-referenced items and increased sensitivity for items encoded in relation to a distant other. The main effect of agent and its interaction with stimulation are illustrated in Figure 1.

**Figure 1.**
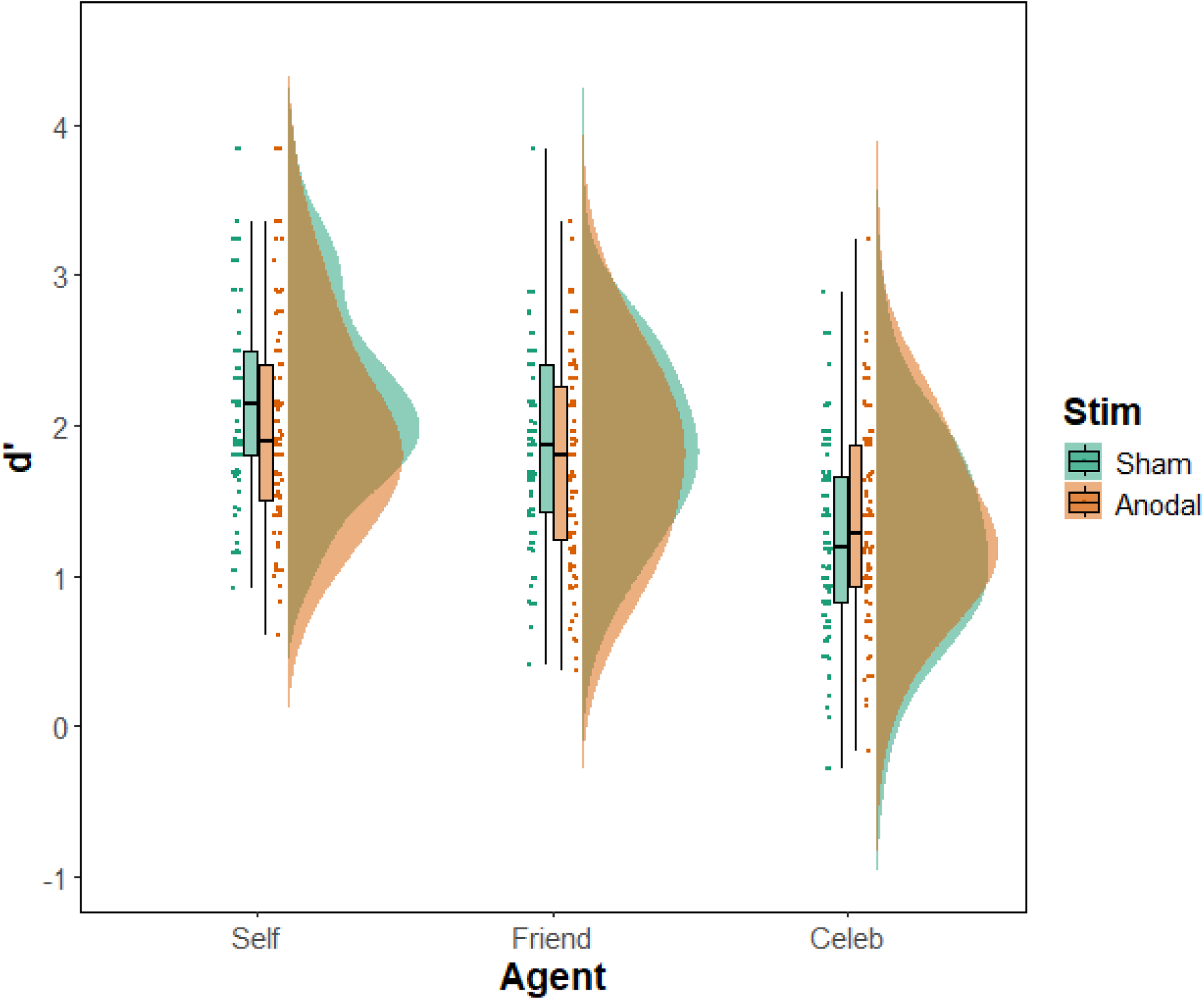
Item Memory Sensitivity (d’) Scores across both Stimulation and Agent Conditions collapsed across stimulation region.

#### Metacognitive Efficiency (meta-d’/d’)

Data from two participants were excluded due to extremely low memory sensitivity (d′ < 0.10). Mauchly’s test indicated violations of the sphericity assumption for several within-subjects effects (all ps < .05). Accordingly, Greenhouse–Geisser corrections were applied to the degrees of freedom and associated p-values where appropriate. Full model outputs are reported in Supplementary Material 2. Below, we focus on the primary results of interest.

A significant main effect of Agent was observed for metacognitive efficiency, F(1.84, 173.02) = 84.27, p < .001, η²G = 0.13, indicating that metacognitive efficiency varied as a function of the encoded agent. Post hoc comparisons showed that self-referenced items were associated with greater metacognitive efficiency than items encoded in relation to a close other, t(97) = 2.55, p = .01, d = 0.18, and a distant other, t(97) = 12.30, p < .001, d = 0.87. In addition, items encoded in relation to a close other showed greater metacognitive efficiency than those encoded in relation to a distant other, t(97) = 9.75, p < .001, d = 0.69.

There was no evidence that stimulation modulated these agent-related differences. Neither the interaction between Agent and Stimulation Type, F(1.84, 173.02) = 0.56, p = .56, η²G = 0.001, nor the three-way interaction between Agent, Stimulation Type, and Region, F(1.84, 173.02) = 0.33, p = .70, η²G = 0.001, reached significance.

No main effect of Valence on metacognitive efficiency was observed, F(1, 94) = 0.53, p = .47, η²G = 0.001. However, there was a significant interaction between Agent and Valence, F(1.89, 177.39) = 24.57, p < .001, η²G = 0.03, indicating that agent-related differences in metacognitive efficiency varied as a function of stimulus valence. Follow-up analyses revealed a substantially larger agent effect for positive words, F(2, 194) = 100.30, p < .001, η²G = 0.22, compared with negative words, F(1.86, 180.43) = 15.23, p < .001, η²G = 0.04.

For positive stimuli, self-referenced items showed greater metacognitive efficiency than items encoded in relation to a close other, t(97) = 4.26, p < .001, d = 0.38, and a distant other, t(97) = 13.83, p < .001, d = 1.25. For negative stimuli, self-referenced items showed greater metacognitive efficiency than items encoded in relation to a distant other, t(97) = 4.55, p < .001, d = 0.44, but did not differ from items encoded in relation to a close other, t(97) = −0.43, p = .67, d = −0.04. Items encoded in relation to a close other showed greater metacognitive efficiency than those encoded in relation to a distant other for negative stimuli, t(97) = 4.98, p < .001, d = 0.48.

No interactions involving Stimulation Type were observed for any valence-dependent effects (all ps between .44 and .91). Together, these results indicate robust agent- and valence-dependent differences in metacognitive efficiency, with no evidence that stimulation to either region modulated these effects.

### Source Memory

Means and standard deviations across all conditions are presented in Table 2.

**Table 2.**
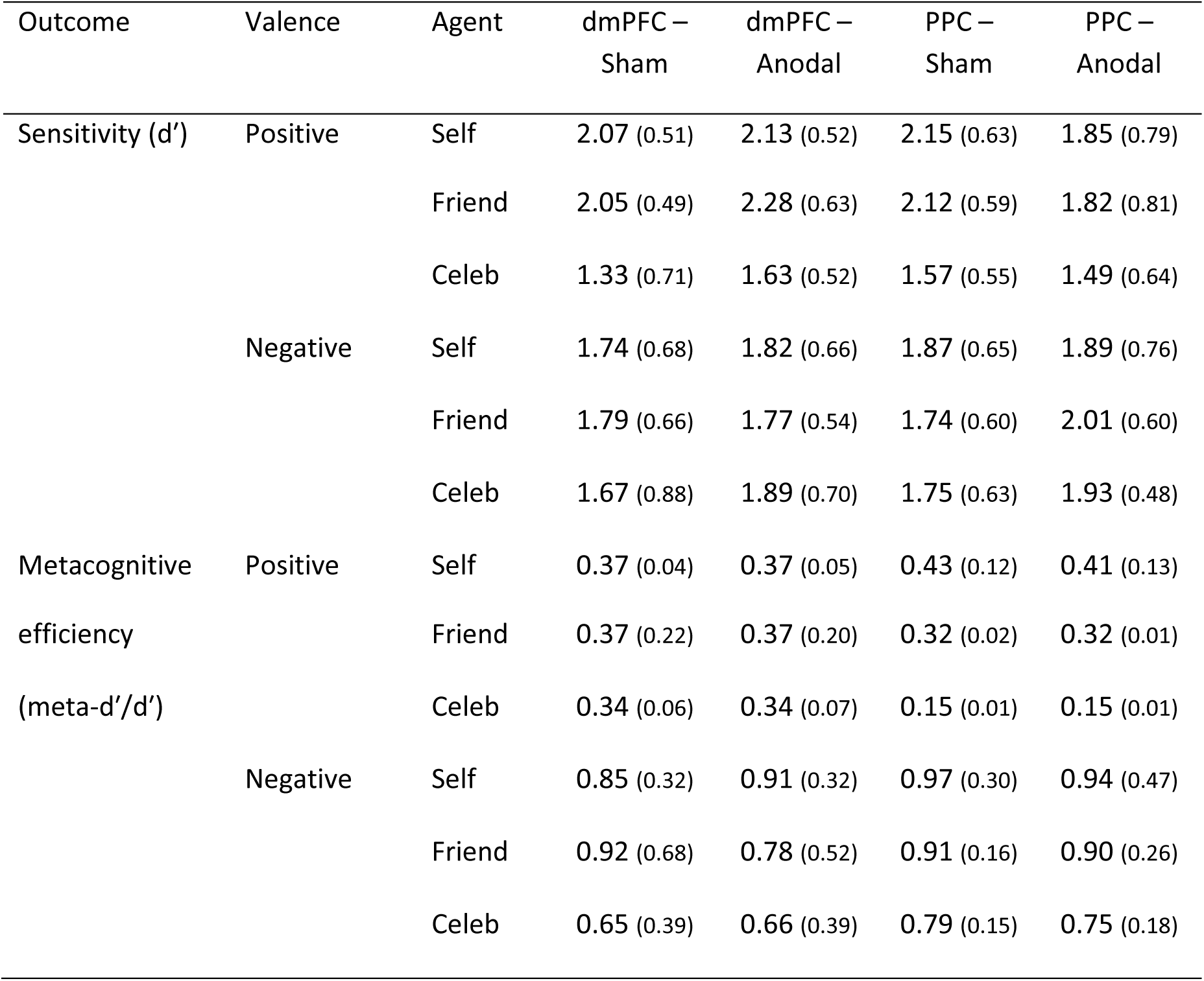
Means and standard deviations across all tasks as per stimulation condition and region of stimulation for source memory.

#### Sensitivity (d’)

A significant main effect of Agent was observed for source memory, F(2, 192) = 20.23, p < .001, η²G = 0.04. Post hoc comparisons indicated that source attribution accuracy differed between items encoded in relation to the self and those encoded in relation to a celebrity, t(197) = 5.45, p < .001, d = 0.44, as well as between items encoded in relation to a close other and those encoded in relation to a celebrity, t(197) = 5.56, p < .001, d = 0.45. In contrast, source attribution accuracy did not differ between items encoded in relation to the self and those encoded in relation to a close other, t(197) = −0.11, p = .91, d = −0.01.

There was also a significant interaction between Agent and Valence, F(2, 192) = 20.67, p < .001, η²G = 0.04, indicating that agent-related differences in source memory varied as a function of emotional valence. Follow-up analyses revealed a robust effect of Agent for positive words, F(2, 198) = 38.63, p < .001, η²G = 0.15. For positive stimuli, source attribution accuracy was higher for self-referenced items than for celebrity-referenced items, t(99) = 7.50, p < .001, d = 0.87, and higher for close-other-referenced items than for celebrity-referenced items, t(99) = 7.72, p < .001, d = 0.89. No difference was observed between self- and close-other-referenced items, t(99) = −0.22, p = .82, d = −0.03.

A significant three-way interaction between Region, Stimulation Type, and Valence was observed, F(1, 96) = 4.78, p = .03, η²G = 0.01. Follow-up analyses conducted separately for each stimulation site revealed that the interaction between Stimulation Type and Valence was not significant for stimulation targeting the dorsomedial prefrontal cortex, F(1, 48) = 0.58, p = .45, η²G = 0.002. In contrast, a significant interaction between Stimulation Type and Valence was observed for stimulation targeting the posterior parietal cortex, F(1, 48) = 5.00, p = .03, η²G = 0.02.

To further characterise this effect, valence-related differences in source memory were examined separately for sham and anodal stimulation of the posterior parietal cortex. Under sham stimulation, source attribution accuracy did not differ significantly between positive and negative items, t(24) = 1.37, p = .18, d = 0.27. Similarly, under anodal stimulation, the difference between positive and negative items did not reach statistical significance, t(24) = −1.78, p = .09, d = −0.36. Complementary analyses examining the effect of stimulation within each valence condition indicated no significant differences in source memory for positive items, t(48) = 1.50, p = .14, d = 0.42, or for negative items, t(48) = −1.14, p = .26, d = −0.32.

Although individual simple effects did not reach significance, the overall interaction pattern indicates that stimulation of the posterior parietal cortex altered the relative balance of source attributions for positive and negative information, shifting from an initial bias favouring positive items towards a relative advantage for negative items (see Figure 2).

**Figure 2.**
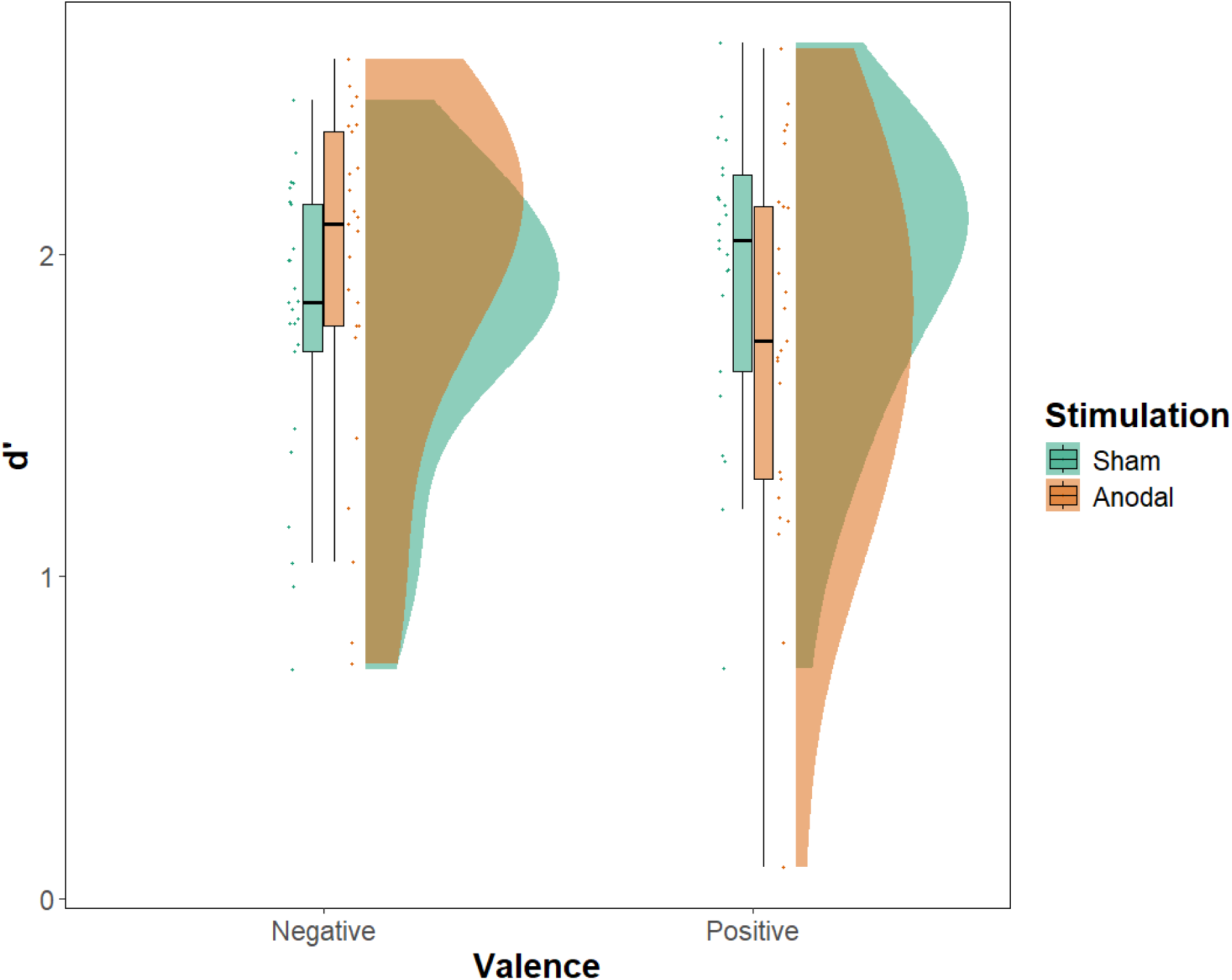
Source memory sensitivity (d′) as a function of stimulation condition and valence following posterior parietal cortex (PPC) stimulation.

#### Metacognitive Efficiency

Data from four participants were excluded due to extremely low memory sensitivity (d′ < 0.10), and a further four participants were excluded due to extreme values exceeding three standard deviations from the condition mean. Mauchly’s test indicated violations of the sphericity assumption for several within-subjects effects (all ps < .05). Accordingly, Greenhouse–Geisser corrections were applied to the degrees of freedom and associated p-values where appropriate. Full model outputs are reported in Supplementary Material (S1). Below, we present the primary results of interest.

A significant main effect of Agent was observed, F(2, 176) = 47.42, p < .001, η²G = 0.07. Post hoc comparisons showed that memory performance differed across all agent conditions. Specifically, self-referenced items were associated with higher performance than items encoded in relation to a close other, t(88) = 2.39, p = .02, d = 0.17, and higher performance than items encoded in relation to a celebrity, t(88) = 9.37, p < .001, d = 0.65. In addition, items encoded in relation to a close other showed higher performance than those encoded in relation to a celebrity, t(88) = 6.98, p < .001, d = 0.49.

A significant main effect of Valence was also observed, F(1, 88) = 277.97, p < .001, η²G = 0.47, indicating substantially higher performance for one valence category relative to the other.

#### Mood change and adverse effects

To assess whether stimulation influenced mood or tolerability, separate two-way ANOVAs were conducted on change scores for negative mood, positive mood, and total adverse effects. Each analysis included Region (PPC vs. dmPFC) and Stimulation (active vs. sham) as between-subjects factors. Means and standard deviations are presented in Table 3.

**Table 3.** Descriptive statistics (mean ± SD) for mood change scores and total adverse effects by stimulation site and condition.

|  | PPC – Sham<br>(n = 25) | PPC – Active<br>(n = 25) | dmPFC – Sham<br>(n = 25) | dmPFC – Active<br>(n = 25) |
| --- | --- | --- | --- | --- |
| Negative mood | -2.64 ± 6.65 | -1.56 ± 5.84 | -0.35 ± 5.63 | -1.28 ± 3.71 |
| Positive mood | -0.30 ± 4.41 | -1.23 ± 2.37 | -0.72 ± 3.27 | -0.58 ± 2.94 |
| Adverse effects | 3.24 ± 3.18 | 2.20 ± 3.15 | 2.56 ± 1.85 | 2.88 ± 2.24 |

*Note.* Mood change scores were calculated as post- minus pre-stimulation composite scores. Negative mood reflects the summed ratings of afraid, confused, sad, angry, tired, and tense, and positive mood reflects the summed ratings of energetic and happy. Total adverse effects scores represent the sum across all adverse effects items (maximum score = 40).

For negative mood change, there was no significant main effect of Region, *F*(1, 96) = 1.33, *p* = .25, η²p = .014, no main effect of Stimulation, *F*(1, 96) = 0.00, *p* = .95, η²p < .001, and no Region × Stimulation interaction, *F*(1, 96) = 0.82, *p* = .37, η²p = .009.

Similarly, for positive mood change, there were no significant main effects of Region, *F*(1, 96) = 0.03, *p* = .86, η²p < .001, or Stimulation, *F*(1, 96) = 0.36, *p* = .55, η²p = .004, and no Region × Stimulation interaction, *F*(1, 96) = 0.65, *p* = .42, η²p = .007.

Analysis of total adverse effects scores revealed no significant main effect of Region, *F*(1, 96) = 0.00, *p* = 1.00, η²p < .001, no main effect of Stimulation, *F*(1, 96) = 0.46, *p* = .50, η²p = .005, and no Region × Stimulation interaction, *F*(1, 96) = 1.62, *p* = .21, η²p = .017.

Together, these analyses indicate that stimulation did not differentially affect mood or adverse effects across regions or stimulation conditions, suggesting that the observed memory effects are unlikely to be attributable to changes in affect or tolerability.

#### Blinding

A chi-square test of independence was conducted to examine whether guessing accuracy differed across the two regions. Accuracy (correct vs. incorrect) was significantly associated with region, χ²(1, *N* = 100) = 7.89, *p* = .005. As shown in Table X, participants produced a higher proportion of correct guesses in Region 1 (34/50; 68%) than in Region 2 (20/50; 40%).

To assess performance relative to chance, binomial tests were conducted separately for each region (chance = 0.5). In Region 1, the proportion of correct guesses was significantly greater than chance (34/50; *p* = .02). In contrast, performance in Region 2 did not differ significantly from chance (20/50; *p* = .89).

## Discussion

The present study investigated how focal tDCS to the dorsomedial prefrontal cortex (dmPFC) and posterior parietal cortex (PPC) modulates self-referential memory across three levels of social distance: self, close other, and celebrity. As expected, we observed a robust self-reference effect, with memory performance highest for self-related information and close-other judgments falling intermediately between self and celebrity. This pattern reinforces the well characterised gradient of mnemonic advantage tied to personal relevance and aligns with previous behavioural work showing graded self, close-other, and familiar-other advantages (Chirtop et al., 2025; Serbun et al., 2011). Notably, focal tDCS to either the dmPFC or the PPC attenuated the SRE in relation to distant others with no effect on memory for close others. Moreover, the present results indicate that anodal stimulation of the PPC exerted valence-specific effects on source memory, selectively improving memory for negative information while impairing memory for positive material across self, friend, and celebrity conditions.

This pattern of stimulation effect offers important insight into how medial prefrontal and posterior parietal regions support the encoding of person information across a gradient of familiarity. Rather than selectively enhancing memory for socially distant targets, stimulation of the dmPFC and PPC attenuated the self-reference effect, indicating a reduction in the mnemonic prioritisation typically afforded to self-related information. Both dmPFC and PPC are implicated in controlled reflective processes, person knowledge retrieval, and attentional allocation, which are typically engaged more strongly for unfamiliar or low-relevance social targets (Ciaramelli et al., 2008, 2020; Sestieri et al., 2017; Wagner et al., 2012). Recent work has extended these accounts by showing that the neural representation of others becomes increasingly differentiated from the self representation as social distance increases, with celebrity targets showing the greatest differentiation (Courtney & Meyer, 2020). The present findings suggest that neuromodulation may act to rebalance representational or attentional weighting across the social mnemonic gradient rather than selectively amplifying processing for distant others. In this context, the observed reduction of the self-reference effect, alongside valence-specific effects of PPC stimulation on source memory, points to a network-level modulation of control and monitoring processes rather than enhanced encoding of any single social category.

The observed pattern may reflect how the social brain differentially supports the construction and updating of person representations across levels of familiarity. The dmPFC is consistently implicated in forming trait impressions, generating mental state inferences, and integrating sparse social information when thinking about others who are not well known to the observer (Baetens et al., 2014; Ferrari et al., 2016). These findings converge with causal evidence from focal high-definition tDCS, which suggests that dmPFC stimulation can alter the integration of other-related information into self-representations during explicit, higher-order social cognitive tasks (Martin, Dzafic, et al., 2017). Rather than selectively enhancing memory for socially distant targets, neuromodulation of the dmPFC in the present study appeared to reduce the relative mnemonic prioritisation of self-related information, consistent with a rebalancing of controlled reflective processes across social targets. Processing socially distant individuals is thought to require greater constructive effort and greater reliance on domain-general reflective systems than processing the self, which draws on well-established personal schemas. Modulating dmPFC activity may therefore alter the relative contribution of these processes without producing net gains in memory for unfamiliar individuals. The parallel effects observed with PPC stimulation further suggest that these operations depend on coordinated interactions between medial and parietal subsystems of the default mode network, which together support attentional control, monitoring, and the elaboration of social information during encoding and retrieval (Li et al., 2014).

These results also align closely with the self–other representational gradient described by Denny et al. (2012), who demonstrated that ventral medial PFC regions support introspective and affectively rich self-knowledge, whereas dorsomedial regions are recruited more strongly when thinking about psychologically distant others. More recent work has refined this account by showing that representational similarity in medial PFC varies continuously with subjective closeness across multiple social categories, ranging from the self to close others, acquaintances, and celebrities (Courtney & Meyer, 2020). The present findings are consistent with this organisational framework. Memory for self-related information, which draws on robust and highly accessible representations, remained largely unaffected by stimulation. Rather than selectively enhancing memory for socially distant targets, tDCS appeared to attenuate the relative advantage typically afforded to self-related material, thereby reducing the self-reference effect. In this context, stimulation may have altered the balance of engagement along the self–other gradient, influencing how cognitive and neural resources were distributed across social targets without producing uniform gains in memory performance.

Finally, these findings contribute to a growing literature examining whether non-invasive brain stimulation can modulate self-referential processing and related social cognitive functions. Although some studies report effects of dmPFC stimulation on self-referential encoding or metacognitive bias (Burden et al., 2021; Mattes & Soutschek, 2025), others find more limited or task-specific effects, particularly when stimuli are neutral or when processing relies on automatic rather than reflective operations (Bao et al., 2021). Our data support the view that stimulation effects are more likely to emerge when tasks engage controlled, effortful social cognitive processes, such as constructing representations of unfamiliar others, rather than when tasks rely on well established and strongly encoded self or close-other schemas. This distinction may help explain the heterogeneity in the stimulation literature and suggests that functional specificity within the social brain offers a principled framework for predicting when neuromodulation will influence social memory.

Although stimulation influenced item memory accuracy, there was no corresponding change in metacognitive efficiency, suggesting that tDCS altered the quality of mnemonic evidence without affecting the ability to monitor that evidence. Both the dorsomedial prefrontal cortex (dmPFC) and the posterior parietal cortex (PPC) are involved in memory performance (Berryhill, 2012; Martin, Dzafic, et al., 2017; Sestieri et al., 2017), yet neither region is believed to provide a general mechanism for metacognitive monitoring in simple item recognition tasks. The PPC contributes to attentional selection and the accumulation of evidence during retrieval, so stimulation can strengthen or weaken the memory signal used for recognition without changing how confidence reflects that signal. In a similar way, mPFC contributions to metacognition appear to be domain specific, supporting self-referential or socially inferential forms of evaluation rather than general confidence judgments for basic episodic decisions (Fleming et al., 2014; Mattes & Soutschek, 2025). When stimulation affects the strength of memory evidence but does not alter monitoring networks in anterior prefrontal, cingulate, or insular regions, accuracy can change while the relation between evidence and confidence remains stable. As a result, metacognitive efficiency is preserved even when stimulation modulates memory performance.

The present results indicate that anodal stimulation of the PPC has valence-specific effects on source memory, improving memory for negative information while impairing memory for positive material across self, friend, and celebrity conditions. This polarity is consistent with proposals that the PPC contributes to selective prioritisation of salient stimuli, particularly those with greater evolutionary or motivational relevance (Klein et al., 2008). Negative words are typically processed with greater attentional weight and stronger contextual encoding (Dent & Martin, 2023; Narhi-Martinez et al., 2023); enhancing PPC excitability may therefore amplify the perceptual–attentional mechanisms that facilitate binding of contextual source features for threat-related information. Conversely, the disruption of positive source memory suggests that increasing PPC excitability biases limited neural resources toward negative information at the expense of more weakly prioritised positive content. Together, these findings suggest that PPC stimulation does not uniformly enhance memory, but instead modulates the emotional weighting of mnemonic processes, favouring better source encoding for negative information while diminishing contextual memory for positive events.

The present findings should be interpreted in light of several limitations. The use of short-term recognition of isolated verbal stimuli may have limited sensitivity to stimulation effects, particularly given that the dmPFC is more strongly engaged during evaluative, contextual, and socially meaningful forms of memory (Amodio & Frith, 2006; Van Overwalle, 2009). Moreover, social memory differs according to characteristics of both close and distant others (Courtney & Meyer, 2020; Dent & Martin, 2023; Kokici et al., 2023; Krienen et al., 2010; Mitchell et al., 2006; Sumar & Martin, 2024; Symons & Johnson, 1997) and should be assessed in future studies. Although focal montages improve spatial precision, current spread beyond the targeted regions cannot be excluded, raising the possibility that effects reflect modulation of broader frontoparietal networks. In addition, tDCS effects are state-dependent and subject to substantial inter-individual variability, which may have contributed to heterogeneous effects in the absence of individualised electric field modelling or physiological measures (Niemann et al., 2024). For example, given documented cultural, sex- and gender-related differences in social cognition, emotional processing, and responsiveness to non-invasive brain stimulation, future studies should explicitly examine whether the observed effects vary accordingly (Adenzato et al., 2017; Martin et al., 2021; Martin, Huang, et al., 2017; Martin, Su, et al., 2019). Finally, the focus on short-term outcomes precludes conclusions about longer-term consolidation processes or underlying network-level mechanisms, which should be addressed in future work using more ecologically valid tasks and multimodal approaches.

In summary, focal stimulation of the PPC and dmPFC produced both shared and dissociable effects on episodic memory. Stimulation to either region attenuated the self-reference effect in relation to distant others. By contrast, PPC stimulation exerted a valence-dependent influence on source memory, strengthening contextual memory for negative stimuli while weakening it for positive stimuli, consistent with differential modulation of affective prioritisation processes. Together, these patterns raise the possibility that focal tDCS engages multiple mechanisms, with item recognition and source memory reflecting partially separable processes that may be differentially sensitive to parietal and medial prefrontal stimulation. Future work combining stimulation with neuroimaging or computational modelling, and extending investigation to clinical or ageing populations, will be important for determining whether these effects reflect common network-level modulation or distinct processes whose functional impact varies across social and emotional contexts.

