## Supplemental Table 1 for "Common and distinct contributions of the dorsomedial prefrontal and posterior parietal cortices to social episodic memory"

**Supplementary Material (S1)**

***Item Memory***

*Sensitivity (d’)*

Repeated-Measures ANOVA Results

| Effect | df | F | p | η²G |
| --- | --- | --- | --- | --- |
| Agent | 2, 192 | 167.61 | < .001 | .213 |
| Agent × Region | 2, 192 | 2.27 | .107 | .004 |
| Agent × Stim | 2, 192 | 6.82 | .001 | .011 |
| Agent × Region × Stim | 2, 192 | 0.62 | .540 | .001 |
| Valence | 1, 96 | 2.71 | .103 | .004 |
| Valence × Region | 1, 96 | 3.15 | .079 | .005 |
| Valence × Stim | 1, 96 | 0.01 | .936 | < .001 |
| Valence × Region × Stim | 1, 96 | 0.01 | .926 | < .001 |
| Agent × Valence | 2, 192 | 0.39 | .680 | .001 |
| Agent × Valence × Region | 2, 192 | 0.99 | .373 | .002 |
| Agent × Valence × Stim | 2, 192 | 0.10 | .905 | < .001 |
| Agent × Valence × Region × Stim | 2, 192 | 0.27 | .767 | < .001 |
| Region | 1, 96 | 0.10 | .755 | < .001 |
| Stim | 1, 96 | 0.37 | .545 | .002 |
| Region × Stim | 1, 96 | 0.06 | .808 | < .001 |

Note. Type III sums of squares were used. η²G = generalized eta squared.

*Metacognitive Sensitivity (meta d’/d’)*

Repeated-Measures ANOVA Results

| Effect | df | F | p | η²G |
| --- | --- | --- | --- | --- |
| Agent | 1.84, 173.02 | 84.27 | < .001 | .128 |
| Agent × Stim | 1.84, 173.02 | 0.56 | .558 | .001 |
| Agent × Region | 1.84, 173.02 | 4.73 | .012 | .008 |
| Agent × Stim × Region | 1.84, 173.02 | 0.33 | .699 | .001 |
| Valence | 1, 94 | 0.53 | .469 | .001 |
| Valence × Stim | 1, 94 | 0.01 | .913 | < .001 |
| Valence × Region | 1, 94 | 21.45 | < .001 | .026 |
| Valence × Stim × Region | 1, 94 | 0.04 | .836 | < .001 |
| Agent × Valence | 1.89, 177.39 | 24.57 | < .001 | .032 |
| Agent × Valence × Stim | 1.89, 177.39 | 0.12 | .880 | < .001 |
| Agent × Valence × Region | 1.89, 177.39 | 2.23 | .113 | .003 |
| Agent × Valence × Stim × Region | 1.89, 177.39 | 0.82 | .438 | .001 |
| Stim | 1, 94 | 2.13 | .148 | .013 |
| Region | 1, 94 | 0.54 | .464 | .003 |
| Stim × Region | 1, 94 | 2.88 | .093 | .018 |

Note. Type III sums of squares were used. Greenhouse–Geisser–corrected degrees of freedom are reported where Mauchly’s test indicated violation of sphericity. η²G = generalized eta squared.

***Source Memory***

*Item Memory (d’)*

Repeated-Measures ANOVA Results

| Effect | df | F | p | η²G |
| --- | --- | --- | --- | --- |
| Agent | 2, 192 | 20.23 | < .001 | .044 |
| Agent × Stim | 2, 192 | 1.67 | .191 | .004 |
| Agent × Region | 2, 192 | 0.48 | .619 | .001 |
| Agent × Stim × Region | 2, 192 | 0.14 | .872 | < .001 |
| Valence | 1, 96 | 0.77 | .382 | .002 |
| Valence × Stim | 1, 96 | 1.42 | .237 | .003 |
| Valence × Region | 1, 96 | 2.26 | .136 | .005 |
| Valence × Stim × Region | 1, 96 | 4.78 | .031 | .010 |
| Agent × Valence | 2, 192 | 20.67 | < .001 | .038 |
| Agent × Valence × Stim | 2, 192 | 0.11 | .897 | < .001 |
| Agent × Valence × Region | 2, 192 | 1.11 | .333 | .002 |
| Agent × Valence × Stim × Region | 2, 192 | 1.06 | .347 | .002 |
| Stim | 1, 96 | 0.49 | .488 | .002 |
| Region | 1, 96 | 0.00 | .970 | < .001 |
| Stim × Region | 1, 96 | 1.18 | .279 | .005 |

Note. Type III sums of squares were used. η²G = generalized eta squared.

*Metacognitive Efficiency (Meta d’/d’)*

Repeated-Measures ANOVA Results

| Effect | df | F | p | η²G |
| --- | --- | --- | --- | --- |
| Agent | 2, 176 | 47.42 | < .001 | .074 |
| Agent × Stim | 2, 176 | 0.70 | .496 | .001 |
| Agent × Region | 2, 176 | 3.32 | .038 | .006 |
| Agent × Stim × Region | 2, 176 | 1.44 | .240 | .002 |
| Valence | 1, 88 | 277.97 | < .001 | .475 |
| Valence × Stim | 1, 88 | 0.11 | .740 | < .001 |
| Valence × Region | 1, 88 | 5.59 | .020 | .018 |
| Valence × Stim × Region | 1, 88 | 0.00 | .987 | < .001 |
| Agent × Valence | 2, 176 | 1.76 | .176 | .003 |
| Agent × Valence × Stim | 2, 176 | 0.76 | .467 | .001 |
| Agent × Valence × Region | 2, 176 | 6.53 | .002 | .012 |
| Agent × Valence × Stim × Region | 2, 176 | 1.04 | .357 | .002 |
| Stim | 1, 88 | 0.18 | .674 | .001 |
| Region | 1, 88 | 0.04 | .841 | < .001 |
| Stim × Region | 1, 88 | 0.01 | .925 | < .001 |

Note. Type III sums of squares were used. η²G = generalized eta squared. Where Mauchly’s test indicated violation of sphericity, corrected statistics were used.
